# Biased neural representation of feature-based attention in the human brain

**DOI:** 10.1101/688226

**Authors:** Mengyuan Gong, Taosheng Liu

## Abstract

Selective attention is a core cognitive function for efficient processing of information. Although it is well known that attention can modulate neural responses in many brain areas, the computational principles underlying attentional modulation remain unclear. Contrary to the prevailing view of a high-dimensional, distributed neural representation, here we show a surprisingly simple, biased neural representation for feature-based attention in a large dataset including five human fMRI studies. We found that when participants selected one feature from a compound stimulus, voxels in many cortical areas responded consistently higher to one attended feature over the other. This univariate bias was robust at the level of single brain areas and consistent across brain areas within individual subjects. Importantly, this univariate bias showed a progressively stronger magnitude along the cortical hierarchy. In frontoparietal areas, the bias was strongest and contributed largely to pattern-based decoding, whereas early visual areas lacked such a bias. These findings suggest a gradual transition from a more analog to a more abstract representation of attentional priority along the cortical hierarchy. Biased neural responses in high-level areas likely reflect a low-dimensional neural code that facilitates robust representation and simple read-out of cognitive variables.

## Introduction

The human brain has the capacity to represent an enormous variety of information. How neural activities represent sensory and cognitive information remains a fundamental question in system and cognitive neuroscience. Answers to this question will inform us how neural processing support adaptive behavior, thus providing important clues regarding the computational principles of the brain.

The prevailing view of neural coding is that the brain uses distributed neural activities to represent information. This view is best illustrated by work that examined neural activities in sensory areas of the brain. Classical neurophysiological studies showed that neurons in early visual areas have smooth tuning functions that span a range of feature values (e.g., Hubel and Wiesel, 1962; Blasdel, 1992; Maunsell, and Van Essen, 1983). Hence, a single stimulus feature would evoke different responses across a population of such neurons, resulting in a specific profile of population activity. Computational studies have demonstrated that such population responses can be used for encoding and decoding sensory information (Pouget, 2000). In parallel to these neuronal level findings, human functional magnetic resonance imaging (fMRI) studies have shown that patterns of BOLD responses can be used to decode and reconstruct visual stimulus (e.g., Kamitani and Tong, 2005; Brouwer and Heeger, 2009; Kay et al., 2008).

Although there is a general consensus that stimulus properties are represented via distributed population activity in sensory areas, much less is known about how cognitive variables are represented in the brain. Cognitive functions related to task control and target selection have been widely associated with activity in parietal and prefrontal cortical areas, collectively known as the Multiple-Demand (MD) network (Duncan, 2010). A key feature of neurons in this network, in contrast to sensory neurons, is that they can flexibly adapt their tuning profile to accommodate task demands (Duncan, 2001; Fedorenko et al., 2013). For instance, neurophysiological studies suggest that population-level neural patterns in prefrontal cortex (PFC) reflect the coding of task rule (Stokes et al., 2013; Freedman, et al., 2001), and the pattern of activity in lateral intraparietal areas (LIP) can encode attentional priority (Bisley and Goldberg, 2003) and learned category (Swaminathan and Freedman, 2012). Contrary to this population-based view, however, a recent neurophysiological study reported evidence supporting a scalar neural code for cognitive variable in LIP (Fitzgerald et al., 2013). These researchers performed detailed analysis of LIP neuronal activity from several experiments in which monkeys performed a variety of cognitive tasks (categorization, associative learning, perceptual decision making). Surprisingly, the majority of recorded neurons from a monkey showed similar response profile, exhibiting largely biased response to one task condition (i.e., higher response to category A than category B). This biased population response implies a low-dimensional, rather than high-dimensional, neural representation for cognitive variables.

The finding of a biased response is surprising (Chaffee, 2013) and thus it is important to know whether the biased representation is restricted to non-human primates or also applies to humans. Many human fMRI studies have decoded cognitive variables using multivariate BOLD response patterns in the MD network, without observing any obvious biases to a particular condition in the univariate response (e.g., Li et al., 2007; Liu et al., 2011; Liu and Hou, 2013; Erez and Duncan, 2015; Bettencourt and Xu, 2016). Although such findings are consistent with a mechanism relying on distributed neural activity, the univariate results were obtained by averaging data across subjects, which could obscure a biased response if the direction of bias varied across subjects. Therefore, investigating the existence of biased neural representation requires us to examine neural responses at the single-subject level. Here, we conducted such analyses on a large fMRI dataset from five attention experiments to examine whether biased neural representation are present in the human brain. Selective attention is a key cognitive function and it is thought to be a core operation that enables complex task control (Duncan 2013). Attentional control is also highly associated with activity in the MD network (Corbetta and Shulman, 2002; Scolari et al., 2015), which includes the intraparietal sulcus, the likely human analogue of monkey LIP. Thus, attentional signals in the brain provide a good test case for possible biased representation of a cognitive variable.

In these experiments, subjects were instructed to attend to one of the two alternative features in a compound stimulus that contained both features, which allowed us to measure the attentional signals throughout the human cortex. We examined potential bias in the neural representation of attentional signals at multiple levels of analysis: single brain area, multiple brain areas within a subject, and multiple subjects at the group level (Fig. 1). Note here that the biased representation describes neural level effects; it does not imply attentional bias at the behavioral level. Our analysis revealed a gradient of increasing bias from sensory to high-level areas, suggesting that low-dimensional neural representations are utilized in the human brain to encode cognitive variables such as attentional priority.

**Fig 1.**
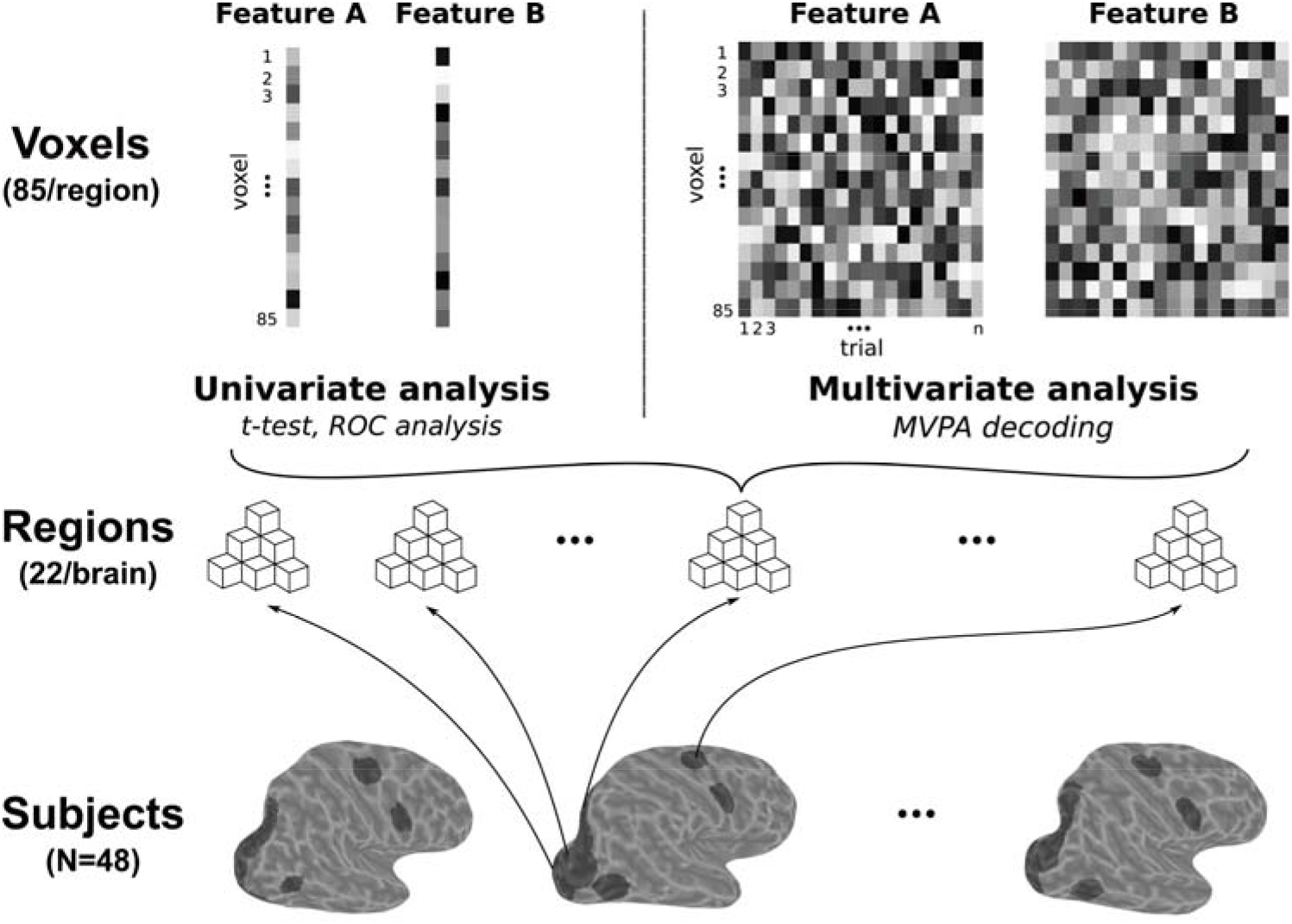
Overview of the data structure and roadmap for the analyses. Bottom row: fMRI data from 48 subjects performing attentional selection tasks were analyzed. Three left hemispheres from different subjects were shown, each overlaid with 11 predefined brain areas (dark shaded areas). Middle row: each brain contained 22 areas (11 per hemisphere), each of which was composed of individual voxels (we used 85 voxels for our main analyses, see *Materials and Methods* for voxel selection and S1 Fig). Top row: data structure for the analyses. For the univariate analysis (left), we obtained two response vectors, one for each attention condition, that contained estimated BOLD response from each voxel for each condition (85 element vector). For multivariate analysis (right), we obtained two matrices of response patterns, each containing BOLD response amplitude from each voxel on each trial in each attention condition (a 85×n matrix, n is the number of trials).

## Results

### Overview of experiments

All experiments utilized a similar paradigm, where subjects were cued to attend to one of the two superimposed stimuli at the same spatial location (Fig 2A & B). To facilitate the presentation of the results, we arbitrarily named the two stimuli as feature A and B (see Fig 2A for details). Subjects’ task was to report brief changes (e.g. luminance or moving speed) contingent on the attended feature (see *Materials and Methods*). This task design kept the physical stimuli constant but varied attentional instruction. Thus, differential neural responses in the two experimental conditions reflect attentional modulation, instead of stimulus-related changes. We show representative results from one of the experiments, where subjects were cued to attend to dots moving in either the upper-left or upper-right direction (Fig 2B). Fig 2C shows the overall brain activation during the task and Fig 2D shows the group-averaged mean time courses of fMRI BOLD response in two representative regions (V1 and FEF). There was no univariate difference between the two attention conditions in average BOLD responses across subjects (Fig 2E). In contrast, the attended feature can be reliably decoded from distributed activity patterns in both visual and frontoparietal areas (Fig 2F). We obtained similar results from the other experiments; details can be found in previous publications (Liu et al., 2011; Liu, 2016; Jigo et al., 2018, Gong and Liu, 2019). In total, the dataset contained BOLD data from 48 subjects, each containing 22 brain areas (11 areas per hemisphere).

**Fig 2.**
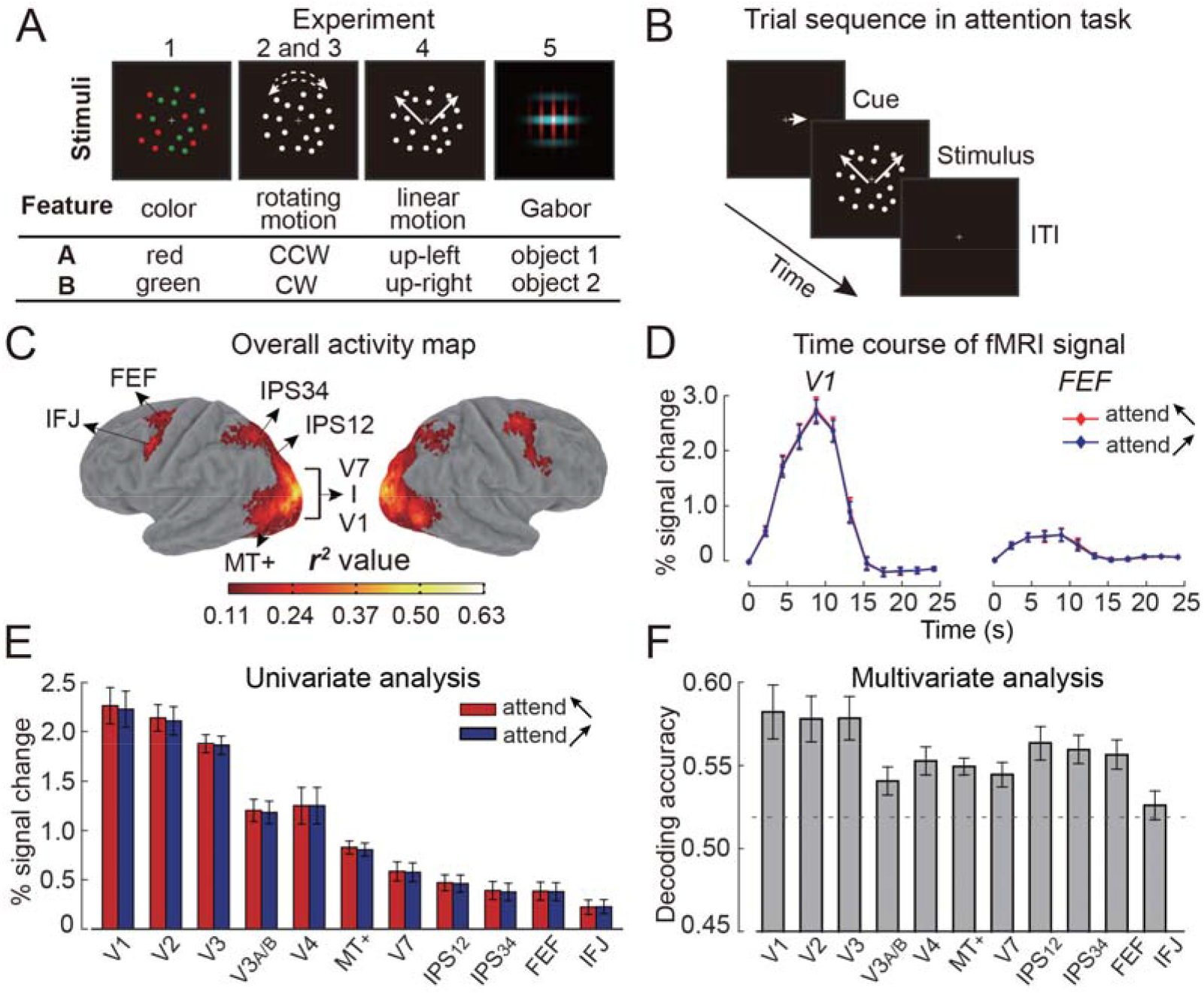
Schematic of stimuli and example results from one experiment. (A) Schematic of stimuli across experiments. Data from five experiments were re-analyzed, with a total of 48 subjects. All experiments employed a compound stimulus containing two features, labeled as ‘A’ and ‘B’, with their meaning explained in the table. (B-F) Example results from Experiment 4 using linear motion (N=12). (B) Trial sequence of the attention task. (C) Overall activity (r^2^) map visualized on an inflated atlas surface. The approximate locations of the key brain areas in occipital and frontoparietal cortex are indicated. IPS: intraparietal sulcus, FEF: frontal eye field, IFJ: inferior frontal junction. (D) Mean fMRI time course for two attention conditions in V1 and FEF. (E) Univariate analysis: mean fMRI responses in visual and frontoparietal areas for each attention condition. (F) Multivariate analysis: decoding accuracy in visual and frontoparietal areas. Gray dashed line indicates maximal significance threshold (corresponding to p<0.05) across brain areas obtained from a permutation test. Error bar denotes standard error of the mean across 12 subjects.

### Biased neural representation of attention across brain areas and subjects

We first examined whether neural activity for the two attention conditions showed mean difference *within each brain area across voxels*, using paired t-tests (see *Materials and Methods*). Fig 3 summarizes all the t-values for all brain areas and subjects, with the color indicating the direction of the bias and the shade indicating its strength. The t-tests were significant in 808 out of 1056 brain areas (~77%) after false discovery rate (FDR) correction for multiple comparisons (corrected p< 0.05, Benjamini and Hochberg, 1995). This is true for both left hemisphere (N=395/528) and right hemisphere (N=413/528), with no obvious difference in the proportion of significant areas between hemispheres (chi-square test for equal proportion: χ^2^=1.71, *p*=0.19). Thus, in many of the brain areas, voxel responses were on average higher for one attended feature than the other, exhibiting a univariate difference, i.e., biased representation.

**Fig 3.**
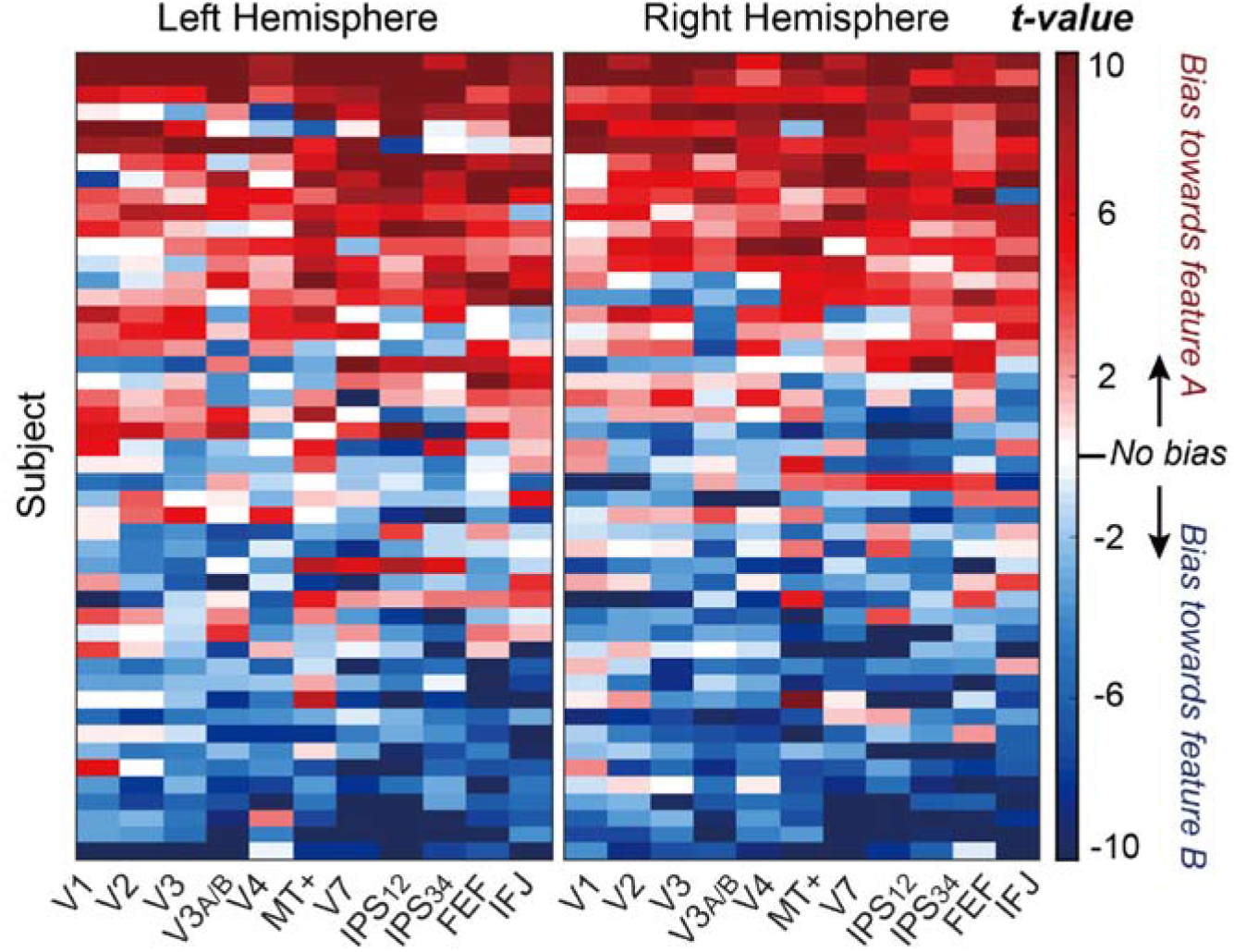
Quantifying the neural bias in individual brain areas and subjects. Data from left and right hemispheres are shown in separate maps. Each row represents data of an individual subject, and each column represents a single brain area. Each cell is color-coded to indicate the *t*-value from the comparison of neural response between two attention conditions (85 voxels were used for this analysis). Red and blue colors indicate the direction of this difference, with their shade indicating the strength of the difference. We arranged this map by sorting the amount of bias per subject (indexed by the mean *t*-values across brain areas), such that from top to bottom, bias progressed from feature A to feature B. For the same data arranged by experiments, see S2 Fig.

Although the direction of the bias varied across subjects, it appeared consistent *within an individual subject across brain areas*. To test the degree of such within-subject consistency, we categorized the direction of bias according to the sign of the t-statistic in each area for each subject (i.e., positive t-values indicated bias towards feature A and negative t-values indicated biased toward feature B). If the direction of such bias is random across brain areas, we should expect approximately half of the brain areas showing a positive bias and the other half showing a negative bias, across the 22 brain areas (11 in each hemisphere). We used a binomial test to evaluate the deviation of observed bias pattern against this null hypothesis and found most of the subjects significantly deviated from the null (N=41/48, maximum *p*=0.0307, *FDR*-corrected). Across subjects, the mean proportion of brain areas that showed the same sign of bias was ~81% (i.e., about 18 out of 22 brain areas showed the same direction of bias).

Thus, although mean BOLD response *across subjects* did not exhibit reliable differences between the two attention conditions (e.g., Fig 2E), voxel responses *within a single brain area* exhibited reliable univariate difference in many of the brain areas examined. The direction of such bias also remained fairly consistent across brain areas within each subject.

### Biased representation increases from sensory to frontoparietal areas

Having observed significant univariate bias, we next quantified such bias to examine how it varies among cortical areas, using a ROC analysis. As an example, Fig 4A shows response distributions from the two attention conditions in a single brain area of a subject. Here the two distributions appear separable with the “attend feature B” condition having larger responses on average than “attend feature A”. This univariate bias can be indexed by the area under the ROC curve (abbreviated as AUC, Fig 4B), which provides a standardized, non-parametric measure of the separation between the two distributions, such that a value of 0.5 indicates no separation and a value of 0 or 1 indicates perfect separation. We calculated AUCs for all 1056 brain areas and found the resulting map of bias very similar to that indexed by t-values (S3 Fig). To quantify the amount of bias regardless of the sign, we rectified the AUC values around 0.5 for individual brain areas (S4 Fig, see *Materials and Methods*). We then averaged AUCs across subjects and performed permutation tests to evaluate their statistical reliability at the group level (Fig 4C, see *Materials and Methods*). We observed statistically significant AUCs in many brain areas (maximal threshold: 0.543), except in early visual areas including V1 and V2 in both hemispheres, and V3 and V4 in the left hemisphere. Furthermore, there is a general trend of increasing AUC values along the cortical hierarchy. This was confirmed by a two-way repeated-measures ANOVA (11 brain areas × 2 hemispheres), showing a strong effect of brain areas (F_(10,470)_=11.29, *p*<0.001, η^2^=.194), without a significant difference between hemispheres (*p*=0.233), or interaction between hemisphere and brain areas (*p*=0.396). In additional analyses, we also explored whether a higher univariate bias is simply due to higher signal-to-noise ratio (SNR) in the fMRI measurement; results showed generally lower SNRs in frontoparietal areas compared to visual areas (S5 Fig).

**Fig 4.**
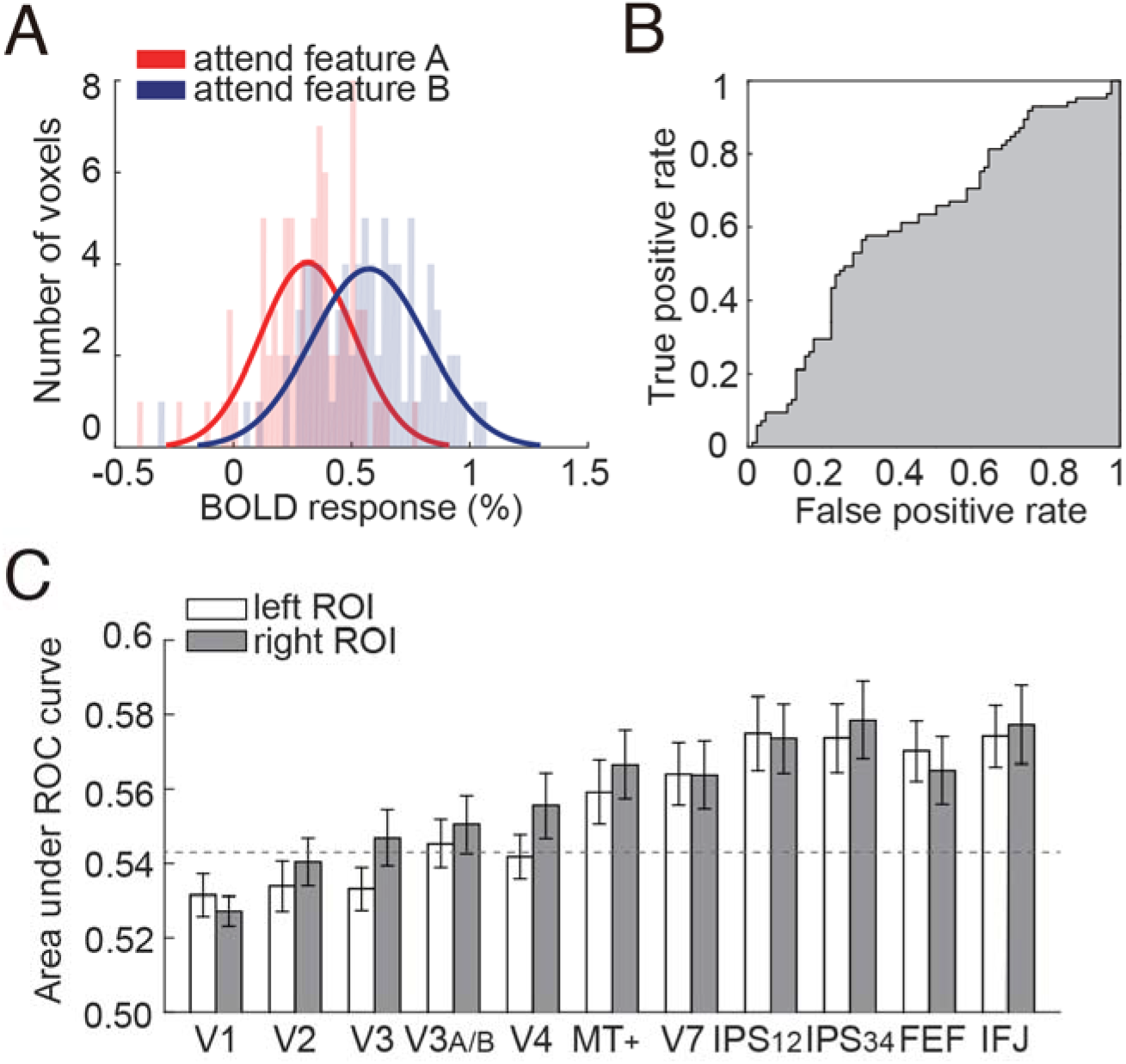
An overview of the ROC analysis. (A) Illustration of the ROC analysis in a single brain area. Histograms show the response distributions (across 85 voxels) in a brain area for the two attention conditions. Solid curves are normal density functions fitted to each distribution. These density curves are for visualization purpose only. (B) The AUC value is calculated from the area under the constructed ROC curve (gray shaded area), a non-parametric measure of the separation between two distributions. (C) Average AUC values across all subjects in all areas of interest. Gray dashed line indicates maximal significance threshold (corresponding to p<0.05) across brain areas obtained from a permutation test.

To further characterize the main effect of brain areas, we grouped the 11 areas into four region groups based on anatomical considerations: V1, extrastriate visual areas (ExS), consisting of V2, V3, V3A/B, V4 and MT+, parietal areas (IPS), consisting of V7/IPS0, IPS12 and IPS34, and prefrontal network (PFC), consisting of FEF and IFJ. Within a region group, we averaged AUCs across constituting brain areas. Because there was no laterality effect, we further collapsed data across two hemispheres. Then, we conducted separate analysis on AUCs to assess whether the biased representation varied with region group and stimulus domain.

#### Comparison across region groups

When grouped into four main anatomical region groups, we observed a clear increase in AUC from V1 to extrastriate visual areas and further into parietal and frontal areas (Fig 5A, red plot). A one-way repeated-measures ANOVA on AUCs revealed a significant effect of region group (F_(3,141)_=23.44, *p*<0.001, η^2^=.333). Pairwise comparisons further showed a stronger bias in frontoparietal areas than that in V1 and ExS (*ps*<0.01), without a significant difference between IPS and PFC (*p*=1.0). To confirm that this result was not due to our specific voxel selection criterion, we repeated the same analysis using different number of voxels (n=105 and 125) and found similar results (Fig 5A, cyan and black plots). A two-way repeated-measures ANOVA (4 region group × 3 voxel number) revealed a main effect of region group (F_(3,282)_=22.12, *p*<0.001, η^2^=.320) and number of voxels (F_(2,282)_=32.15, *p*<0.001, η^2^=.406), as well as a two-factor interaction (F_(6,282)_=6.35, *p*<0.001, η^2^=.119), showing that greater bias was found with fewer voxels in ExS, IPS and PFC (*ps*<0.001), but not in V1 (p=0.559). Because we selected voxels based on the rank-ordered degree of activity during the task, this finding suggests that bias is stronger in more active voxels. Importantly, these results demonstrate the robustness of biased representation of attention in higher-order areas.

**Fig 5.**
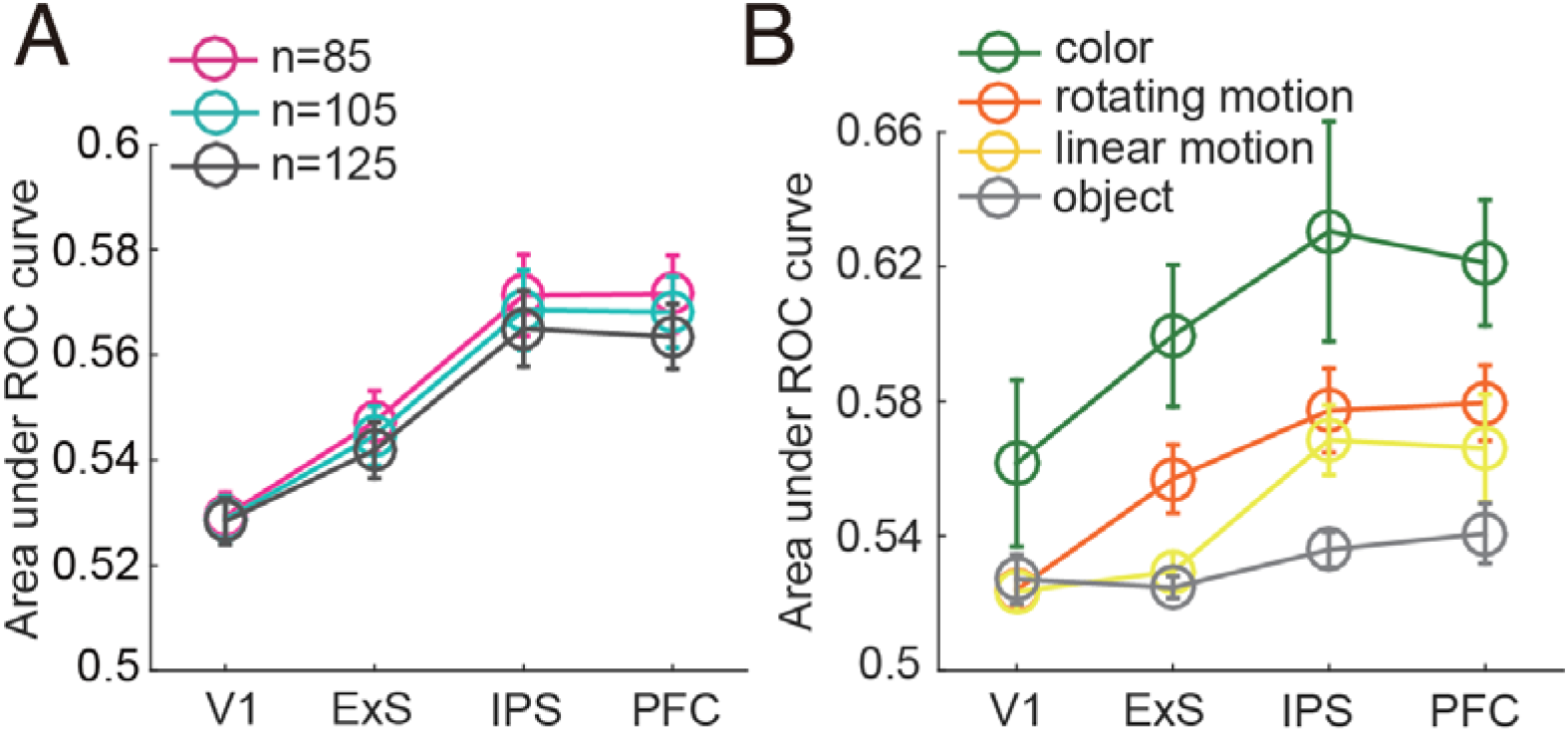
Results from the ROC analysis with varying numbers of included voxels and stimulus domain. (A) Average AUC values for different region groups, obtained with different numbers of voxels (n=85, 105 and 125) in the ROC analysis. (B) Average AUC values for each stimulus domain, obtained with data from different experiments. Error bar denotes standard error of the mean.

#### Comparison across stimulus domains

Because we used different stimuli (i.e., colors, rotating motion, linear motion directions and dynamic objects) across experiments, the amount of bias may differ across stimulus domains. When we plotted the AUCs separately for each stimulus domain (Fig 5B), we observed largest AUCs for colors, intermediate AUCs for motion directions, and smallest AUCs for the dynamic objects. This observation was confirmed by a mixed-effect ANOVA (4 region group × 4 stimuli), which showed a main effect of region group (F_(3,132)_=22.56, *p*<0.001, η^2^=.339) and stimuli (F_(3,44)_=7.28, *p*<0.001, η^2^=.332), as well as a marginally significant two-factor interaction (F_(9,132)=_1.95, *p*=0.051, η^2^=.117). These results indicated that the biased representation of attentional signal was modulated by stimuli, with a decrease of bias along with increasing complexity of the attended information (e.g., from simple features to complex objects). To examine whether the overall pattern of results was mainly contributed by participants from one experiment, we constructed four reduced datasets by removing data from each of the four stimulus domains and conducted four separate ANOVAs on the reduced datasets. All ANOVAs showed a significant main effect of region group (maximum p<0.001), suggesting that the observed increasing bias along the cortical hierarchy is robust and not contributed by data from only one stimulus domain.

### Bias removal produces dissociable effects in sensory and frontoparietal areas

The AUC analysis demonstrates a univariate bias between two attention conditions in many brain areas. Previously we have shown significant above-chance multivariate decoding using pattern classification techniques in all those areas (e.g., Fig 2F). Given that both methods index the neural discriminability between conditions, this raises the question of how much the univariate bias contributes to the multivariate decoding. We used the grand-mean of BOLD signal from each attention condition as a proxy measure of this bias (equivalent to the means of each distribution in Fig 4A). We then subtracted this grand-mean from each attention condition and performed both the ROC analysis and MVPA decoding, separately for each brain area (see *Materials and Methods*). As expected, AUCs in all region groups fell below the significance threshold (i.e., not different from chance) because mean removal essentially eliminated univariate difference (Fig 6A). If multivariate decoding relies mostly on univariate differences, we would expect that removing the mean diminishes the decoding accuracy. Alternatively, if multivariate decoding relies on multi-dimensional pattern variability, we would expect little impact of mean removal on decoding accuracy. We found that mean removal had progressively stronger impact on decoding accuracy along the cortical hierarchy (Fig 6B, for results from individual subjects see S6 Fig). This was confirmed by a two-way repeated-measures ANOVA (region group × mean removal), showing a main effect of mean removal (F_(1,141)_=15.04, *p*<0.001, η^2^=.242), and importantly, a significant interaction between region group and mean removal (F_(3,141)_=3.84, *p*=0.011, η^2^=.076). Follow-up t-tests showed that mean removal produced a significant drop of MVPA-based decoding in ExS (*p*=0.016), IPS (*p*<0.01), and PFC (*p*<0.001), but not in V1 (*p*=0.26). A significant interaction was also obtained if we excluded V1 data from the analysis (F_(2,94)_=6.27, *p*<0.01, η^2^=.118), indicating a relatively larger drop in decoding accuracy due to mean removal in IPS and PFC compared to ExS. Indeed, after mean removal, decoding accuracy in PFC dropped to the chance level (below the gray dashed line in Fig 6B). These results suggest that the progressively stronger univariate bias along the cortical hierarchy, as shown by the ROC analysis, contributes significantly to MVPA-based decoding. Indeed, in the PFC, MVPA decoding appears to rely exclusively on the univariate bias. We obtained similar pattern of results when using different number of voxels (S7 Fig).

**Fig 6.**
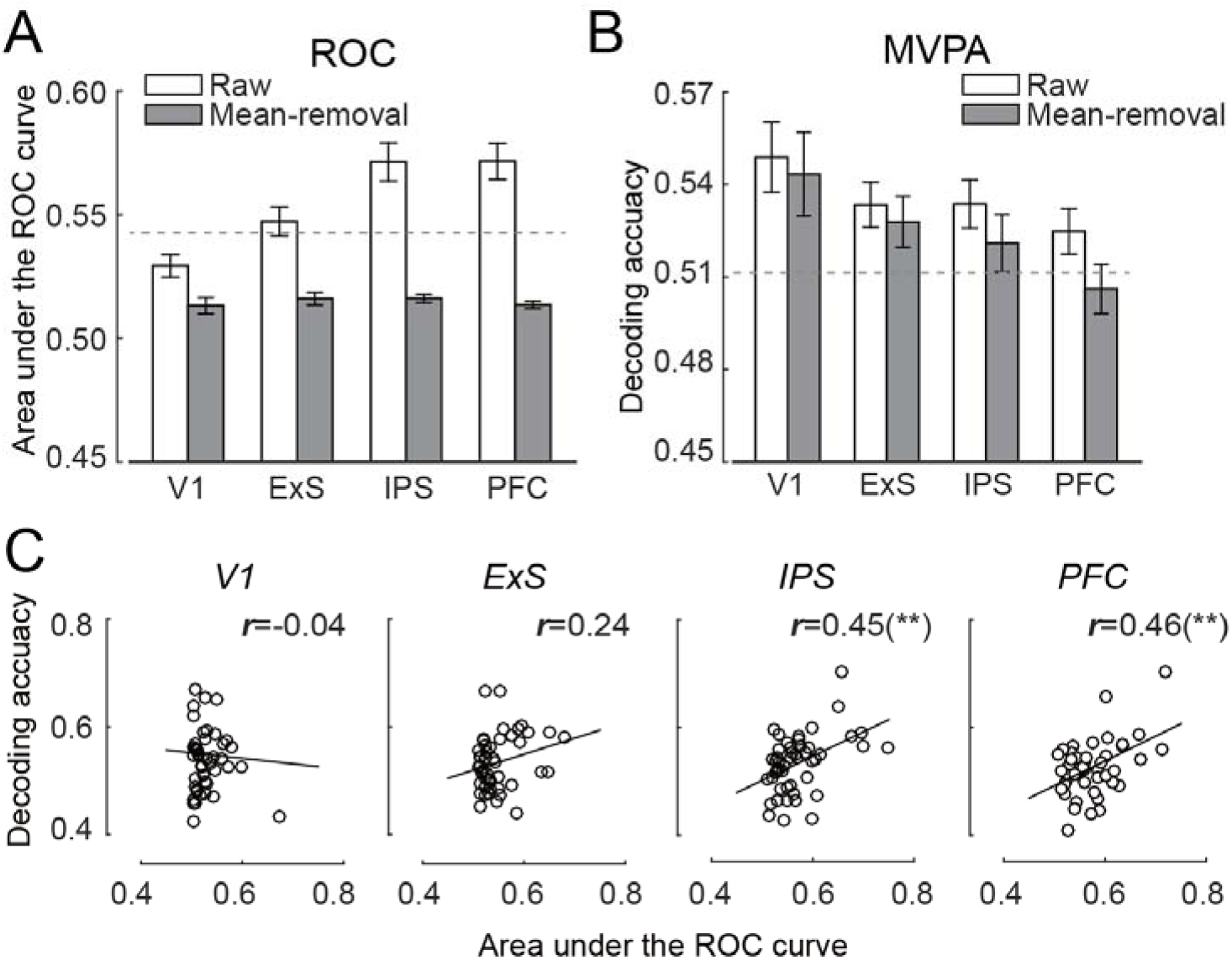
The relationship between ROC and multivariate pattern analysis. (A) ROC results using the original data (Raw) and after removing the grand mean from each attention condition (Mean-removal). (B) MVPA results before and after removing the grand mean from each attention condition. Gray dashed lines indicate maximal significance thresholds (p<0.05) obtained from permutation tests. Error bar denotes SEMs. (C) Inter-subject correlation between AUC and MVPA decoding accuracy for each region group (** indicates statistically significant correlation at *p*<0.01).

To further examine the contribution of univariate bias to MVPA-based decoding, we performed a correlation analysis between AUCs and MVPA decoding accuracies across subjects (Fig. 6C). In particular, for regions where the univariate bias contributes to MVPA-based decoding, we should expect a dependency between these two measures. Indeed, we found significant correlations between these AUC and MVPA decoding in frontoparietal areas (IPS: *r*=0.45, *p*=0.001; PFC: *r*=0.46, *p*=0.001), but no such correlations in sensory areas (V1: *r*=−0.04, *p*=0.78; ExS: *r*=0.24, *p*=0.10). This result further supports the possibility that attention decoding in the frontoparietal network were driven by the univariate bias to a large extent.

### Biased coding does not correlate with behavioral selection

We examined if the observed neural bias was a consequence of preferential behavioral selection. For example, a neural bias in favor of a particular feature may result from a stronger top-down attention to that feature, contributing to higher accuracy and faster reaction time, even though subjects were always instructed to attend equally to individual features. Such a preferential selection should lead to better behavioral performance in the attention tasks. We thus tested this possibility by examining neural and behavior relationship, by correlating the difference in behavioral performance between the two attention conditions (i.e., feature A – feature B) and AUCs. For this analysis, we used raw AUCs without rectification, which captured the direction of the neural bias (Fig 7, see *Materials and Methods).* This analysis revealed no such correlations in any of the region groups between the magnitude of the neural bias and behavioral preference (accuracy: *ps*>0.20; reaction time: *ps*>0.28), making it unlikely that the observed neural biased is due to behavioral preferences.

**Fig 7.**
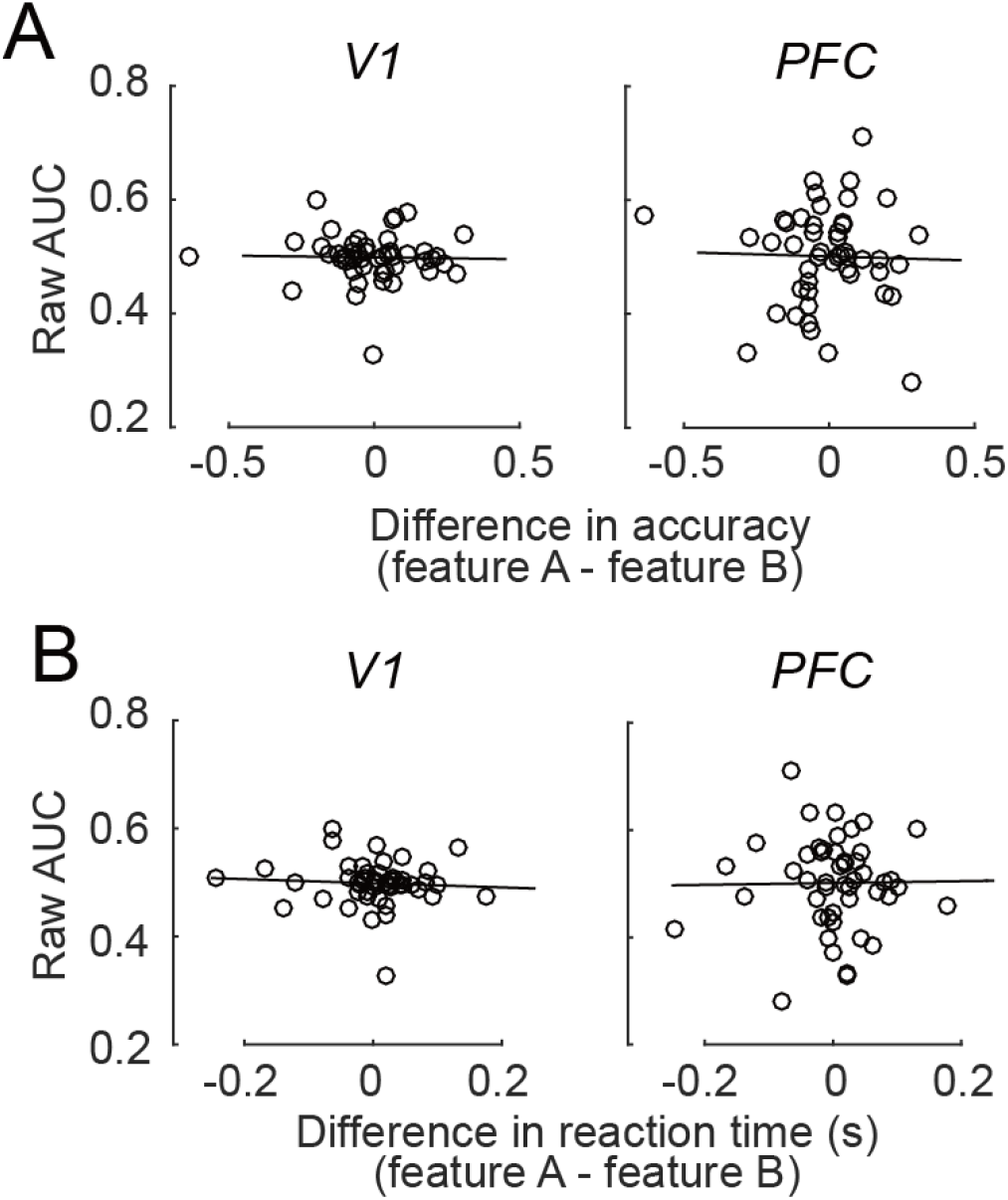
The relationship between raw AUC and behavioral difference. Correlations between raw AUC values and behavioral difference, as indexed by (A) accuracy and (B) reaction time, between two attention conditions in two representative areas (V1 and PFC).

### Equivalent distribution of biased coding at the group level

Lastly, we examined the group-level distribution of the direction (or sign) of the biased representation by assigning each subject a preferred feature (see *Materials and Methods*). We grouped individuals according to the stimuli they viewed during the task (i.e., rotating motion, linear motion, and dynamic object). We excluded data for the attend-color experiment because of the small sample size in that experiment (N=6). Fig 8 shows approximately equal distribution of biased direction across subjects (Chi-square test against equal proportion: *ps*>0.41). Thus, about half of the subjects showed a biased response to feature A and the other half showed the opposite bias. This observation explains the lack of systematic univariate bias at the group level because averaging the two opposite biases likely cancelled each other out.

**Fig 8.**
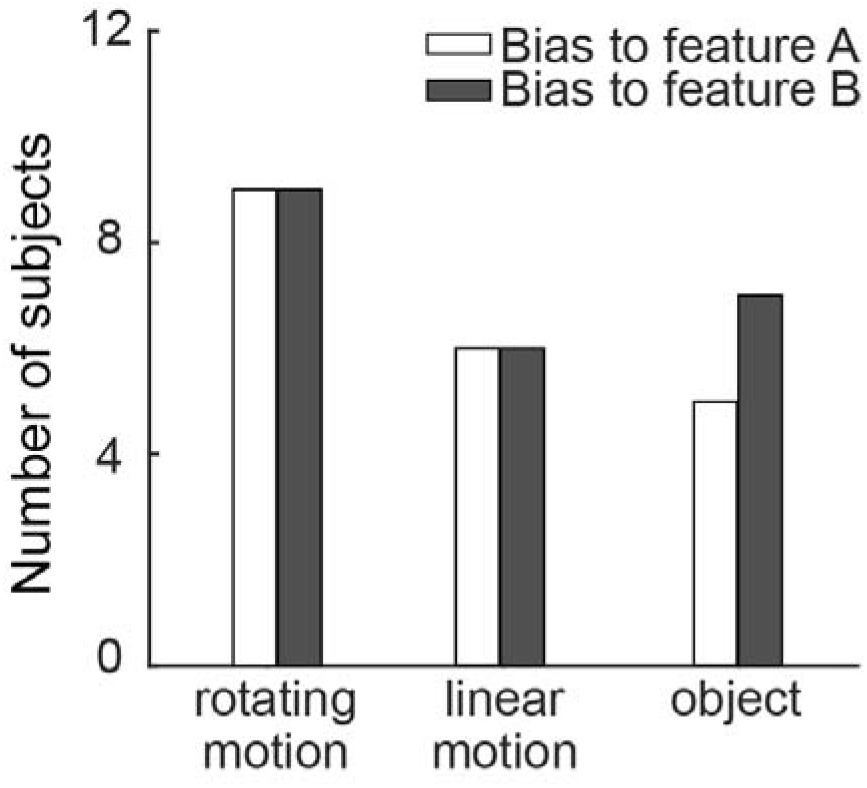
Group-level distribution of biased representation across subjects. The distribution of biased representation across subjects was shown separately for different stimuli (i.e., rotating motion, linear motion, and dynamic object).

## Discussion

We observed a biased neural representation of the attended feature in a large fMRI dataset containing 48 subjects across multiple experiments. This biased representation was robust, as demonstrated by a convergence of findings from single brain area and consistency across multiple brain areas within a subject. At the level of brain areas, we found that voxels in many brain areas showed differential univariate activity between attention conditions. Within individual subjects, the direction of bias remained consistent among brain areas. However, at the group level, the direction of bias varied across subjects with no predominant direction, which also explains the lack of group-average univariate difference in the standard analyses. Importantly, when we quantified the amount of univariate bias using the ROC analysis, we found a progressively stronger biased response from sensory to frontoparietal areas. We ruled out the possibility that the results were due to specific voxel selection criterion, stimulus domain or behavioral preference. Collectively, our findings provide the first evidence for a biased neural representation of cognitive variables in the human brain.

The initial observation of biased representation (Fitzgerald et al., 2013) in non-human primates was indeed rather surprising (Chafee, 2013). However, it is unknown whether such bias is only present in non-human primates who typically undergo extensive training in cognitive tasks before neural recording experiments, or the specific brain area examined, namely LIP. Although our attention task and the tasks performed by the monkeys are different, they shared some formal similarities. First, all tasks were designed to isolate neural signals for cognitive, associative representations without contributions from motor and response-related processes. Second, these cognitive representations establish an association between visual input and a categorical structure as specified by the task. Our results thus extend Fitzgerald et al.’s results to a new class of tasks and to humans with much less training. Here we further leveraged the large sample size and whole-brain coverage afforded by fMRI and show that bias occurs in a multitude of brain areas with consistent direction amongst brain areas at the individual subject level, but the direction of bias varies across subjects. We note that the magnitude of the observed bias here is weaker than the neuronal level data, as Fitzgerald et al., observed nearly all recorded neurons had similar direction of bias. There are likely multiple reasons for this difference in magnitude, such as differences in species, stimulus/task, measured signals (neural spiking vs. BOLD response), and selection criteria (feature selective neurons vs. task-responsive voxels regardless of feature selectivity). Notwithstanding these methodological differences, the findings from both studies converge in demonstrating a biased neural representation for cognitive variables.

Notably, with the whole-brain coverage of fMRI data, we found a dissociable pattern of bias between early visual areas and frontoparietal areas. First, the magnitude of bias grows progressively larger from early visual areas to frontoparietal areas, with an absence of bias in early visual areas. Second, the MVPA analysis on mean removed data suggests that the univariate bias made progressively larger contribution to pattern-based decoding along the cortical hierarchy. In particular, the chance-level decoding in PFC stands in sharp contrast to the unchanged decoding accuracy in V1 after removing the univariate bias (Fig 6B). The contribution of the univariate bias to pattern-based decoding was further supported by the finding of significant correlation between AUC values and decoding accuracies in frontoparietal areas. These contrasting results thus support a distinction between two mechanisms: distributed population representation in sensory areas and biased representation in high-level areas.

From a computational point of view, our results likely reflect different dimensionality of the neural signals at different levels of cortical hierarchy. Specifically, the differential impacts of removing the grand mean on multivariate decoding suggests that sensory areas contain high-dimensional neural signals whereas frontoparietal areas contain low-dimensional, possibly one-dimensional, neural signals. Such a coding scheme is also consistent with general theories of visual information processing, which typically assume that sensory input is processed along hierarchical stages that start with analog representations and gradually transition to task-related, abstract representations (Riesenhuber and Poggio, 1999; Hochstein and Ahissar, 2002; Deco and Rolls, 2004). This transition likely involves changes in the coding properties in different brain areas and our observation of different amount of bias (and signal dimensionality) among cortical areas could be one manifestation of this transition.

Analog representations are naturally implemented with sensory neurons that are continuously tuned to stimulus features (Hubel and Wiesel, 1962; Blasdel, 1992; Maunsell, and Van Essen, 1983). Consistent with this idea, we observed a distributed population response in sensory areas without appreciable univariate bias. Our results are also consistent with a large body of fMRI studies that have reliably decoded the attended stimulus features (Kamitani and Tong, 2005; 2006; Liu et al., 2011; Liu and Hou, 2013) and memorized stimulus features (Serences et al., 2009; Harrison and Tong, 2009; Riggall and Postle, 2012) from activity patterns in early visual areas. Together, these findings support the distributed population representation of cognitive variables in early visual areas.

While analog representation via distributed population response in early sensory areas is well established, how information transitions to an abstract representation in high-level brain areas is much less known. In our data, we found evidence for a biased, or low-dimensional, representation for attended feature in frontal and parietal areas, which are the likely sources of attentional control (Bisley and Goldberg, 2010; Kastner and Ungerleider, 2000) and also part of the Multiple-Demand (MD) network (Duncan, 2013; Fedorenko et al., 2013). It may seem counterintuitive that with the myriad of neurons and their complex connections, the brain uses a low-dimensional, or possibly scalar, code to represent an abstract cognitive variable. Computational analyses, however, have pointed out some benefits of using such a simple code. Because individual neurons are always noisy, a biased response pattern would allow a simple operation, such as averaging, to achieve a reliable representation that is robust to noisy fluctuations in neural systems (see Fitzgerald et al., 2013). Furthermore, such a simple neural code will also simplify the readout of downstream areas and the control of behavior. A related idea was proposed in a modeling study (Ganguli et al., 2008), in which LIP neuronal data from categorization and decision making tasks were found to obey one-dimensional dynamics, such that slowly evolving activity patterns are proportional to spontaneous activity. The investigators suggested that by reducing local neural signals to one-dimensional activity, the brain achieves robust temporal control of behavior such as the timing in shifting attention and crossing a decision threshold during evidence accumulation. Although these proposed benefits of low-dimensional representations reflect different aspects of information coding, they are similar in that unreliable and heterogeneous neural activities from individual neurons can be pooled to achieve more robust representation of cognitive variables.

A natural question concerns how low-dimensional neural activity is generated in the brain. While simulations with simple network models show that local, sparse, recurrent excitatory connections can generate low-dimensional neural activity, it is also possible that coupling among cortical areas plays a role, especially if recurrent activity is weak in individual areas (Ganguli, et al., 2008). A limitation of previous single-unit work is that all the data come from a single brain area, namely LIP. Thus, it is unknown whether low-dimensional neural activity is restricted to one, or a few, brain areas, or is instead a network phenomenon. Our data showed a biased response pattern in the wide-spread MD network, and critically, the direction of such bias was consistent across nodes in this network. Our results thus suggest that network-level interaction could contribute to the generation and maintenance of low-dimensional neural activity.

It is worthwhile considering the generality of biased neural representation. Given our human fMRI data and previous monkey single-unit data were obtained from a variety of behavioral paradigms, biased representation of cognitive variables appears to be a general principle of neural coding in high-level cortical areas. However, we should note that a commonality shared among these behavioral paradigms is that all tasks entail a few (often two) discrete task conditions. It is possible that biased representation is particularly useful in this type of regimes, but would be less useful with more complex task contexts (e.g., an attention task with increased number of features). Theoretical studies suggest that neural dimensionality could scale with task complexity (Fusi, Miller, & Rigotti, 2016; Gao and Ganguli, 2015). This idea has found some support in studies where the number of task conditions appears to drive estimates of dimensionality in monkey PFC (Brincat, et al., 2018; Rigotti et al., 2013). There is also some hint in our data supporting this notion, as the univariate bias was weaker for dynamic multi-feature objects than single features (Fig 5B). Future studies are necessary to systematically evaluate the influence of task and stimulus complexity on the dimensionality of neural signals in the association cortex.

We also do not know how biased representation arises in the first place. Each subject in our dataset only performed a task once in a single scanning session, which does not allow us to test if the direction of the bias persists over a longer period of time. It is possible that the direction of such bias is determined by each individual’s past experience, hence more or less fixed for that individual, or alternatively, such bias arises stochastically when performing a particular task. It would be interesting to examine the consistency and origin of the neural bias in future studies. In addition, because fMRI provides only an indirect and coarse measure of neural activity, future studies that more directly assess neuronal activities (such as single- or multi-unit recordings) would be valuable to provide converging evidence and further insights on the nature of the biased neural representation.

In conclusion, we highlight biased neural representation as a potential mechanism for coding cognitive variables in the brain. Although the simplicity of this coding scheme seems counterintuitive, it can facilitate a robust representation and simple read-out of information critical for stimulus selection and cognitive control. Together with the findings of distributed population representation in early sensory areas, our results suggest a gradual transition from high- to low-dimensional representation along the cortical hierarchy. Such a gradient of neural representation could enable information processing at multiple levels of abstraction to support adaptive behavior.

## Materials and Methods

### Ethics statement

All participants gave informed consent according to the study protocol approved by the Institutional Review Board at Michigan State University (LEGACY08-211).

### Participants

In total, forty-eight subjects (21 females, mean age 25.1 years) from Michigan State University were included across five experiments. We based our sample size on previous studies using similar attention tasks, details of which can be found in previous publications. All had normal or corrected-to-normal vision. Participants were paid for their participation at $20/hr.

### Overview of the experimental procedures

We re-analyzed data from five previously published fMRI experiments (Liu et al., 2011; Liu, 2016; Jigo et al., 2018; Gong and Liu, 2019). The details of the methods can be found in previous publications, so only an abbreviated description is provided here. All experiments used a similar task design: two features were presented in the same location and subjects were cued to attend to one of the features on a trial-by-trial basis. The stimulus features varied across experiments, and we will refer them as feature A and feature B in this report. There are thus two experimental conditions: attend A and attend B. The exact features are as follows. In the first experiment, six subjects attended to a color in a superimposed red-green color display (Liu et al., 2011). In the second and third experiment, six and twelve subjects attended to a motion direction in a superimposed clockwise-counterclockwise rotating dot display (Liu et al., 2011; Jigo et al., 2018). In the fourth experiment, twelve subjects attended to a linear motion direction in a superimposed up-left/up-right moving dot field (Gong and Liu, 2019). In the fifth experiment (Liu, 2016), twelve subjects attended to a dynamic object in a superimposed display containing two Gabor patches (Object 1 or Object 2) that continuously changed their features in multiple dimensions (color, orientation, and spatial frequency). Data for each subject were collected in a single 1.5 – 2 hr scanning session. In total, the dataset contained 48 subjects. The number of trials in each condition (attend A or attend B) varied from 29 to 136 across experiments. The order of conditions was fully randomized.

In all experiments, subjects performed a threshold-level change detection task on the attended feature (e.g., detecting a speedup event in the attended motion direction). Each subject was extensively trained on the task before the scanning session with their performance calibrated by a psychophysical staircase procedure. The task was sufficiently challenging to engage feature selection. In all experiments, we verified that performance did not differ between attend feature A and attend feature B conditions (for details see previous publications).

### Retinotopic mapping

For each subject in each experiment, we ran a separate scanning session of visual field mapping to define visual and parietal topographic areas. We used standard phase-encoded checkerboard stimuli to define retinotopic visual areas (Sereno et al. 1995; DeYoe et al. 1996; Engel et al. 1997) and a memory delay saccade task to map topographic areas in the parietal cortex (Sereno et al. 2001; Schluppeck et al. 2006; Konen and Kastner 2008). All areas were defined and visualized on computationally flattened representations of the cortical surface, which were generated from high-resolution anatomical images using FreeSurfer (http://surfer.nmr.mgh.harvard.edu) and custom Matlab code. Detailed descriptions of the mapping procedure can be found in our previous publications. The following regions of interest (ROIs) in each hemisphere were identified with this procedure: V1, V2, V3, V3A/B, V4, V7, MT+, IPS1 to IPS4.

### Univariate analysis: deconvolution

We used the deconvolution approach by fitting each voxel’s time series with a general linear model whose regressors modeled the two attention conditions with finite impulse responses. The design matrix was pseudo-inversed and multiplied by the time series to obtain an estimate of the hemodynamic response (HRF) evoked by each condition. For each voxel, we computed a goodness of fit measure (r^2^ value), corresponding to the amount of variance explained by the deconvolution model (Gardner et al. 2005). The r^2^ value represents the degree to which the voxel’s response over time is correlated with the attention task. Thus, when using the r^2^ value to select voxels (see below), we essentially selected voxels based on their overall modulation in BOLD response during the task, regardless of any *differential* activity among conditions.

We also performed permutation test to assess the statistical significance of the r^2^ values to aid our voxel selection. For each subject, we repeated the deconvolution analysis for 1000 times, each time with a random re-shuffling of the trial labels. For each of the 1000 analysis, we took the maximum r^2^ value across all voxels in the brain to obtain a distribution of 1000 maximum r^2^ values. This null distribution thus contained the maximum possible r^2^ value expected by chance for all voxels and can be used to assess the statistical significance of the observed r^2^ values while controlling for family-wise Type I error (Nichols and Holmes, 2001). The p-value of each voxel was calculated as the percentile of voxels from the null distribution that exceeded the observed r^2^ value. We also used r^2^ value in conjunction with the anatomical constraints to define two frontal areas as clusters of active voxels during the attention task: frontal eye field (FEF) in the vicinity of the precentral sulcus and superior frontal sulcus, and inferior frontal junction (IFJ) at the intersection between inferior frontal sulcus and inferior portion of precentral sulcus.

### Voxel selection and response calculation

For each ROI, we first eliminated noisy voxels defined as any voxel that showed more than 10% signal change. We then sorted the voxels by their r^2^ value in a descending order and selected the top 85 voxels for each ROI and subject for further analysis. This number of voxels was found in >95% of all ROIs (1056 in total). For ROIs that had fewer than 85 voxels using this criterion, we used all voxels that satisfied the criterion (average: 70 ± 12 voxels). This number of voxels was also approximately the number of active voxels in frontoparietal areas as assessed by the permutation test above (see S1 Fig). Due to the anatomical difference, the proportion of selected voxels was smaller in visual areas than that in frontoparietal areas, we thus repeated the main analyses using r^2^ sorted voxels at 105 and 125 voxels, which included ~90% and ~85% of all ROIs that met this criterion, respectively.

Taking into account the difference in study design (block- vs. event-related) across experiments, we calculated the voxel response by averaging different time windows: 5 – 20 s after trial onset for block-design in Exp. 1 and 2; 4.4 - 11 s after trial onset for event-related design in Exp. 3 to 5. For univariate analysis, we averaged time points from the deconvolved response. For multivariate analysis, we averaged time points from the raw time series for each trial.

### Receiver operating characteristic (ROC) analysis

For each subject and each ROI, we performed a receiver operating characteristic (ROC) analysis on the voxel-wise deconvolved response to quantify the univariate discriminant information between the two attention conditions. The ROC analysis is essentially a classification analysis based on one-dimensional, univariate response. The area under the ROC curve (abbreviated as AUC) indicates the reliability with which an ideal observer could distinguish the attended feature given the response distributions for both conditions (e.g., Britten et al., 1992). In general, we rectified the raw AUC values around 0.5 (e.g., an AUC of 0.45 is rectified to 0.55) to quantify the discriminant information regardless of the direction of bias. For ease of explanation, we refer to the rectified AUC values simply as AUC, and the original, un-rectified AUC as raw AUC. Because the AUC values were always greater than 0.5 (theoretical chance level), we assessed the statistical significance of AUC using a permutation test. We shuffled the labels of trials to obtain an AUC of the shuffled data and repeated this procedure 1000 times to obtain a null distribution of 1000 AUCs for each subject and each ROI. To compute the group-level significance, we concatenated the null distribution across 48 subjects to obtain a 48×1000 matrix and then averaged the values over subjects to obtain a group-level null distribution of 1000 AUC values for each brain area. The 95 percentile of this group-level distribution was thus defined as the statistical significance level (corresponding to p=0.05) to determine if an observed AUC value significantly exceeded the chance level. For the correlation analysis between behavioral difference and AUCs (Fig 7), we used the raw AUCs such that the direction of neural bias (A>B vs. A<B) was represented by values greater or less than 0.5, respectively. Finally, we used the median AUC across ROIs to label each subject’s direction of bias (values greater than 0.5 indicated bias for A, and vice versa). This subject level assignment was then used to assess the distribution of bias at the group level.

### Multivariate pattern analysis (MVPA)

For each voxel and each ROI, we obtained single-trial fMRI response amplitude (see *Voxel selection and response calculation*). We thus obtained a m×n instance matrix for each ROI and attention condition, where m was number of trials and n was number of voxels. We then performed MVPA to discriminate between the two conditions using Fisher linear discriminant analysis. As we often have fewer trials than voxels, which made the estimated covariance matrix non-invertible, we added a ridge coefficient to the diagonal elements of the covariance matrix (Warton, 2008). We performed leave-one-run-out cross-validation to evaluate the classification accuracy, by dividing the data set into test (one run) and training data (remaining runs). This procedure was repeated until each run was tested once. Classification accuracy was averages across folds for each ROI. To assess the contribution of biased coding to MVPA, we subtracted the grand mean of each m×n instance matrix from the instance matrix itself (i.e., mean removal separately for each attention condition) before applying the same MVPA analyses as before. Similar to the permutation test used for ROC analysis, we assessed the significance of decoding accuracy by shuffling the trial labels in the training data and calculated the decoding accuracy on the test data. We repeated this procedure for 1000 times to compute a null distribution for each subject and each ROI. We then averaged the null distributions over all subjects to obtain a group-level distribution of decoding accuracies for each ROI. The 95 percentile of this group-level distribution was thus defined as the statistical significance level (corresponding to p=0.05) to determine if an observed MVPA decoding accuracy significantly exceeded the chance level. In all analyses where multiple statistical tests were conducted, we corrected the p-values using the false discovery rate method (Benjamini and Hochberg, 1995).

## Acknowledgements

We thank Dr. David Zhu and Ms. Scarlett Doyle for their assistance in collecting the neuroimaging data. We also thank Michael Jigo for assistance in data collection and analysis. This work was supported by a NIH grant (R01EY022727).

