## Supplementary materials for "Biased neural representation of feature-based attention in the human brain"

**Figure captions for supplementary materials**

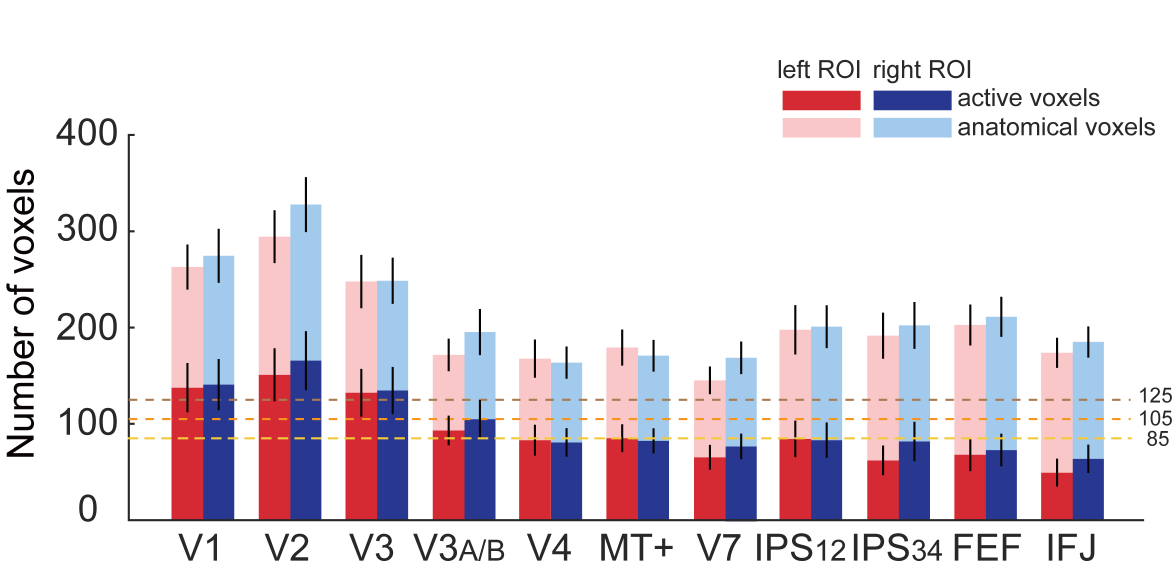

**S1 Fig. Number of anatomical voxels and task-related active voxels**

We assessed each voxel’s responsiveness to the task using a permutation test, which allowed us to derive a p-value for the observed r^2^ value in each voxel (see Materials and Methods). We labeled voxels with p<0.05 as active and plotted the average number of active voxels (dark bars) and the total number of anatomical voxels (light bars) for each pre-defined brain area. Three different levels of voxel inclusion criteria (85, 105, 125) are indicated by horizontal dashed lines. Black vertical lines denote 95% confidence intervals across subjects.

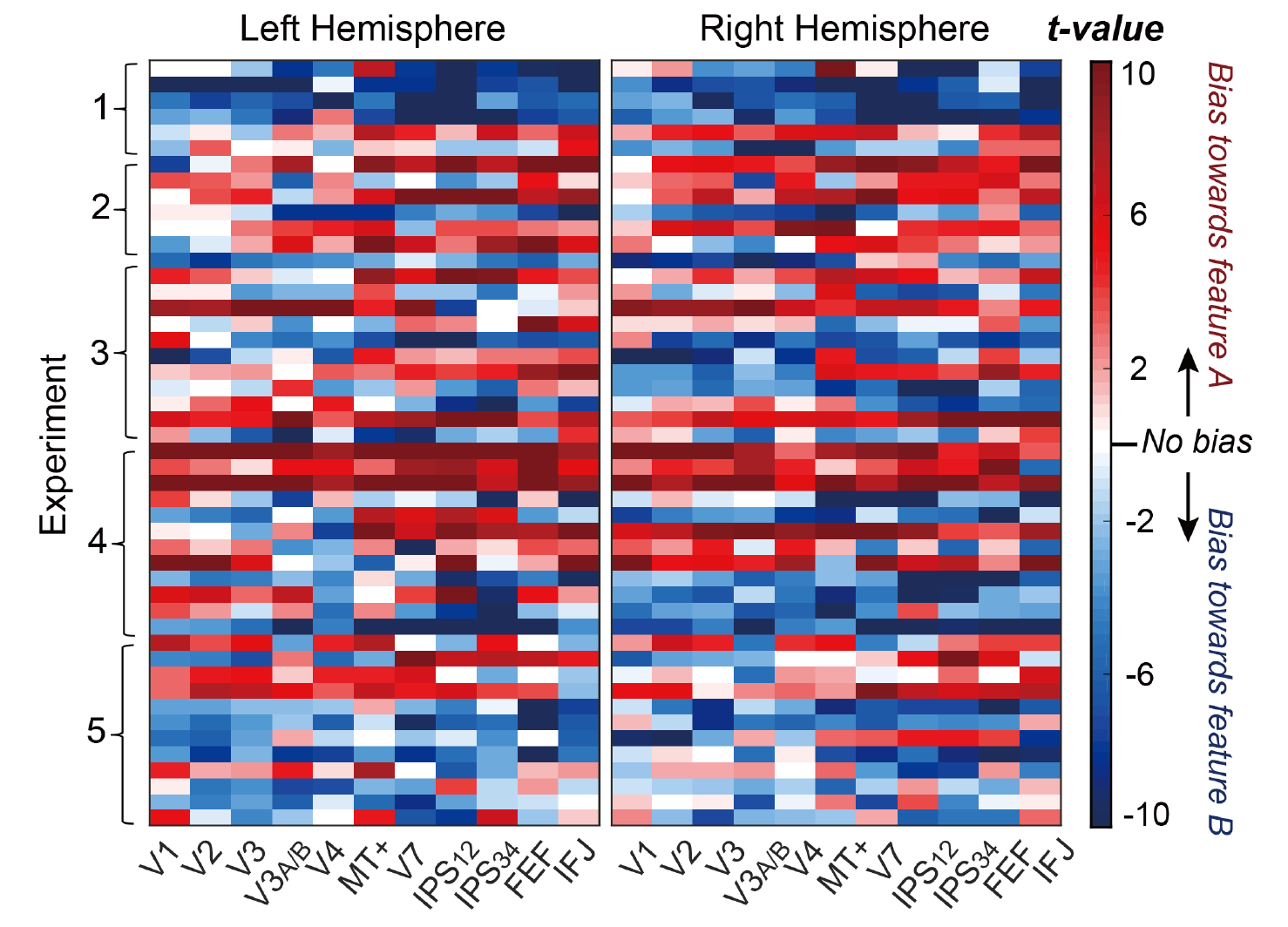

**S2 Fig. Neural bias indexed by t-tests in individual brain areas and subjects.**

This figure shows the same results as in Fig 3, except that the map was arranged by the experiment, as indicated by the left column.

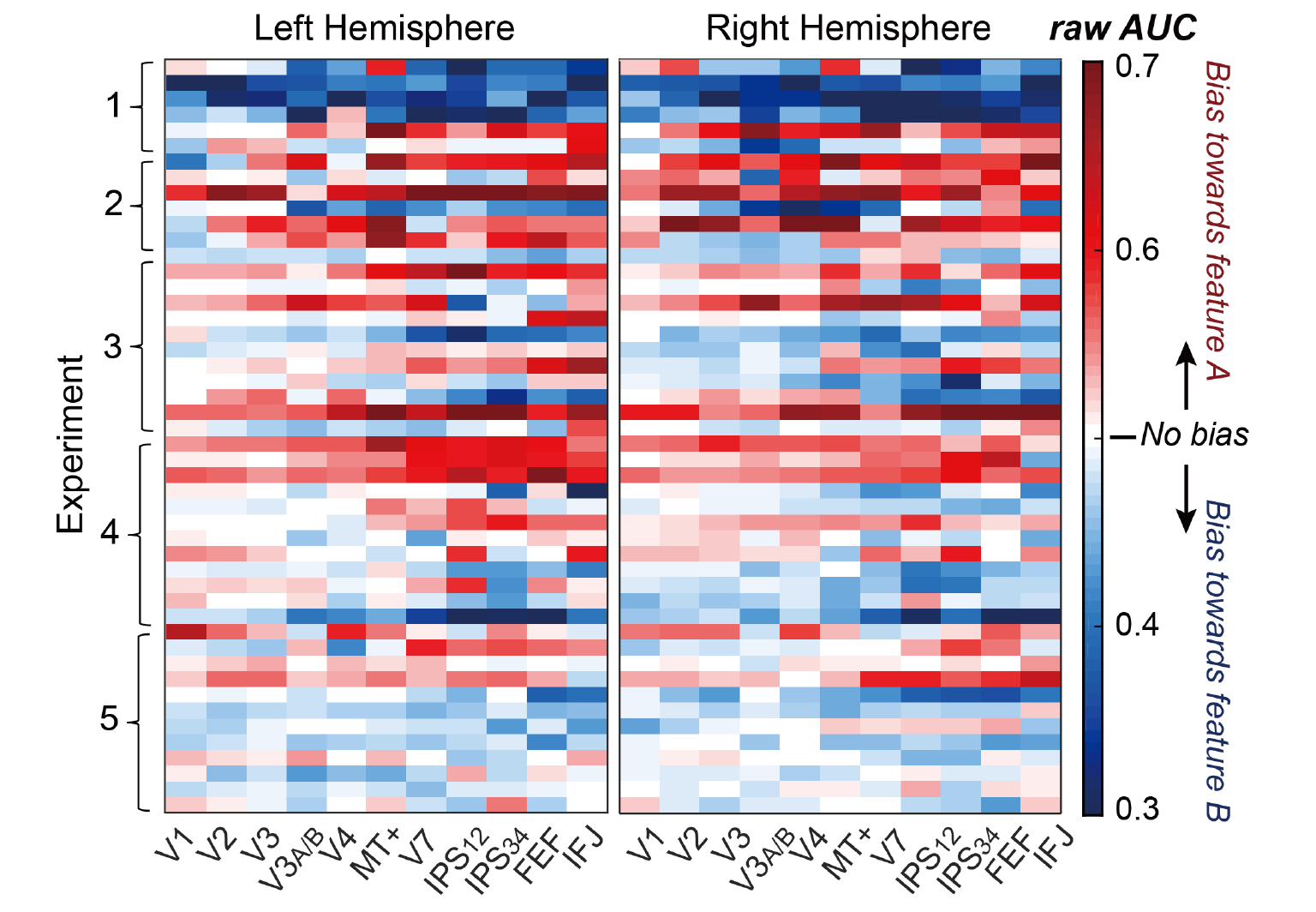

**S3 Fig. Neural bias indexed by raw AUC in individual brain areas and subjects.**

We quantified the biased representation using ROC analysis. Similar to S2 Fig, each cell is color-coded to indicate the raw AUC value from the comparison of neural response between two attention conditions (85 voxels were used for this analysis). Red and blue colors indicate the direction of this difference, with their shade indicating the strength of the difference. The map is arranged by experiments, like S2 above. The sign of the bias in the two maps (S2 and S3) is consistent in more than 95% of the areas.

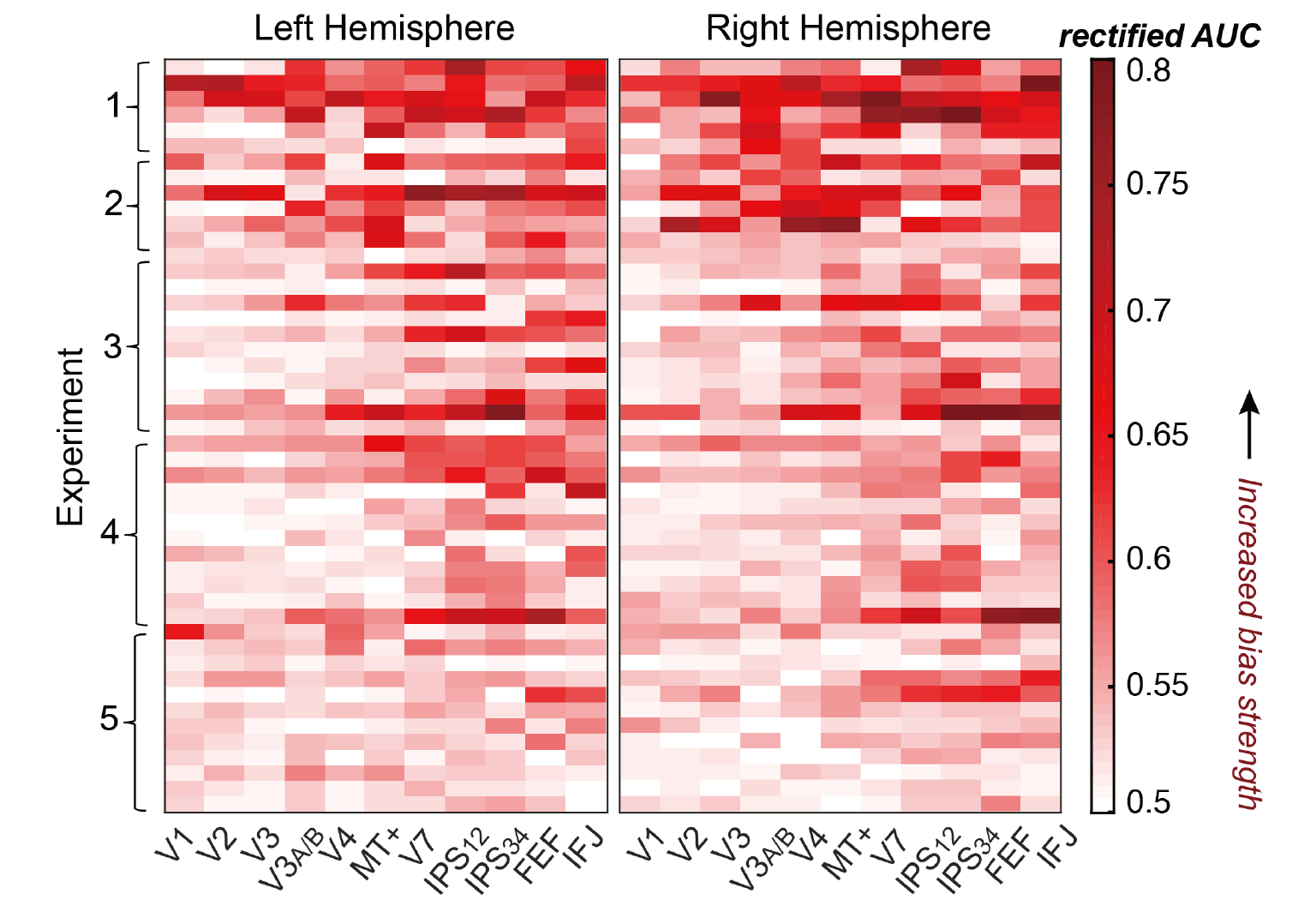

**S4 Fig. Neural bias indexed by rectified AUC in individual brain areas and subjects.**

To show the amount of neural bias in individual subjects, we plotted the rectified AUC in a similar format as S3 Fig. Each cell is coded to indicate the absolute amount of neural bias (rectified AUC value) from the comparison of neural response between two attention conditions. The map is arranged by experiments, like S2 and S3 above.

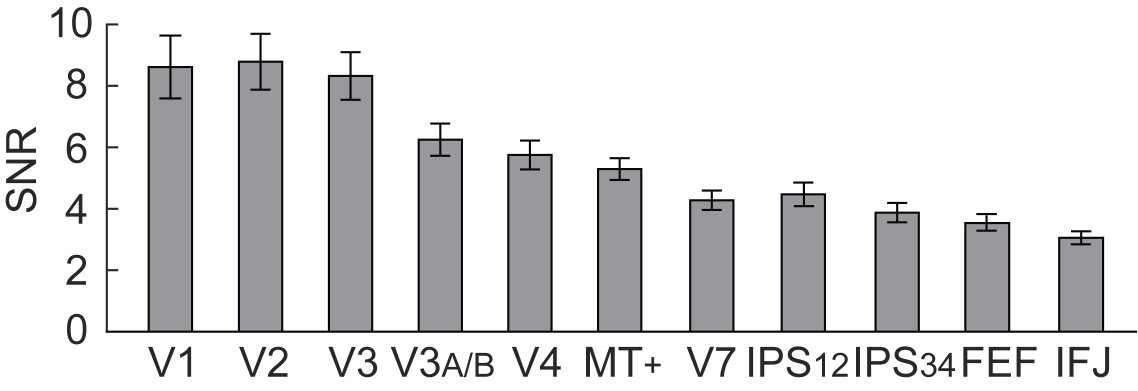

**S5 Fig. Signal-to-noise ratios (SNR) across brain areas.**

Because a higher SNR might lead to a larger AUC value, we assessed whether the SNR in our fMRI data also increased along the cortical hierarchy. For each voxel at each time point and condition, we defined the SNR as the deconvolved BOLD response divided by the error of the deconvolution model (an index of trial-by-trial fluctuations of the BOLD response due to noise). To quantify the overall SNR in each brain area, we averaged the SNR across time points, voxels, conditions and experiments. The averaged time windows were identical to that used for ROC analysis. Contrary to the above expectation, overall SNR decreased from sensory to frontoparietal areas, suggesting the increased AUC values in the latter areas reflect differences in neural representation, but not differences in SNRs. Error bar denotes SEMs.

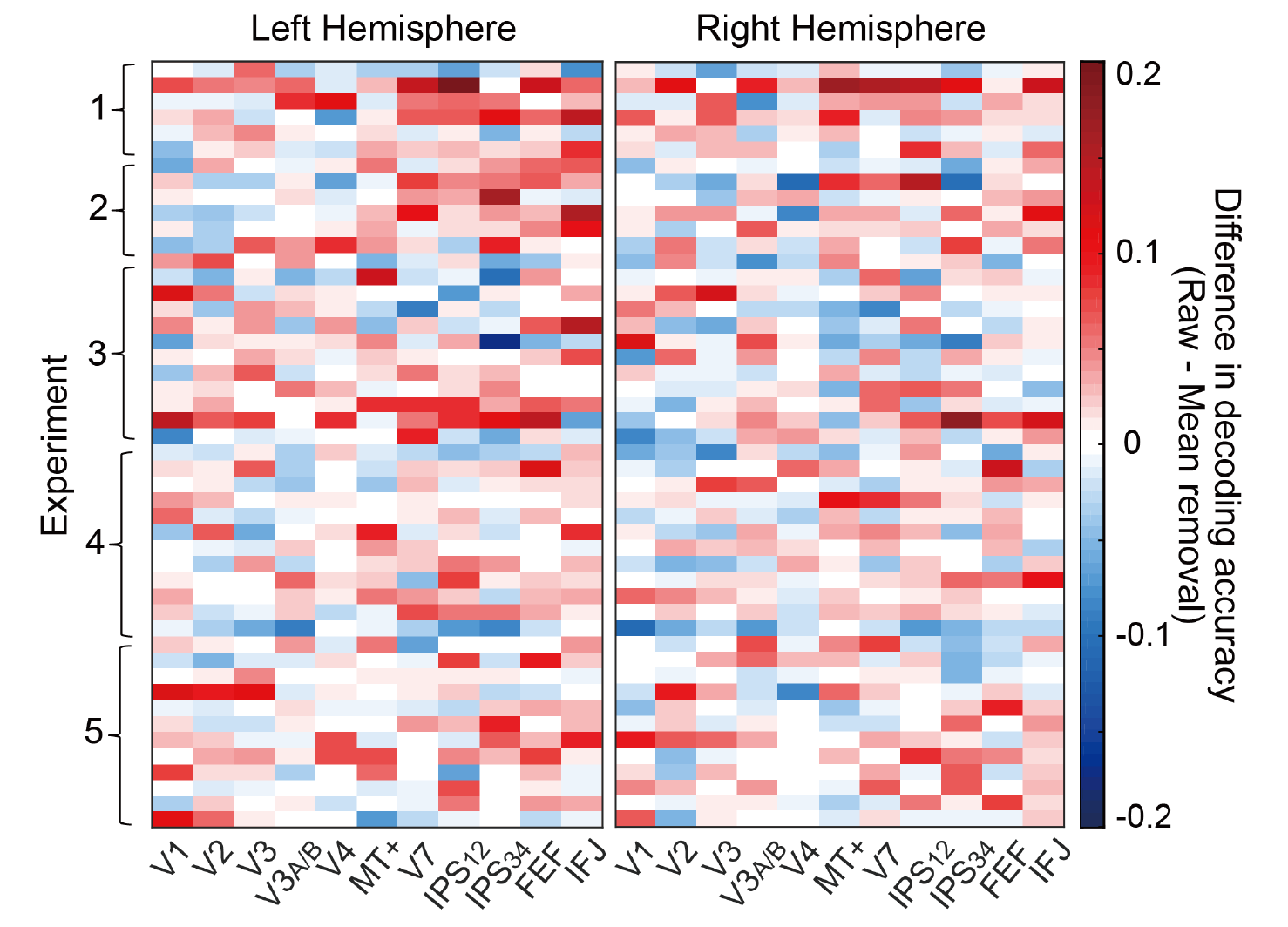

**S6 Fig. The effect of mean removal on MVPA in individual brain areas and subjects.**

This figure shows the difference in MVPA decoding accuracy (Raw – Mean removal) in individual subjects and brain areas, in a similar format as S2 to S4. Positive values indicate a decrease in decoding accuracy due to mean removal; negative values indicate an increase. Overall, frontoparietal areas exhibited a larger decrease in decoding accuracy than visual areas due to of mean removal.

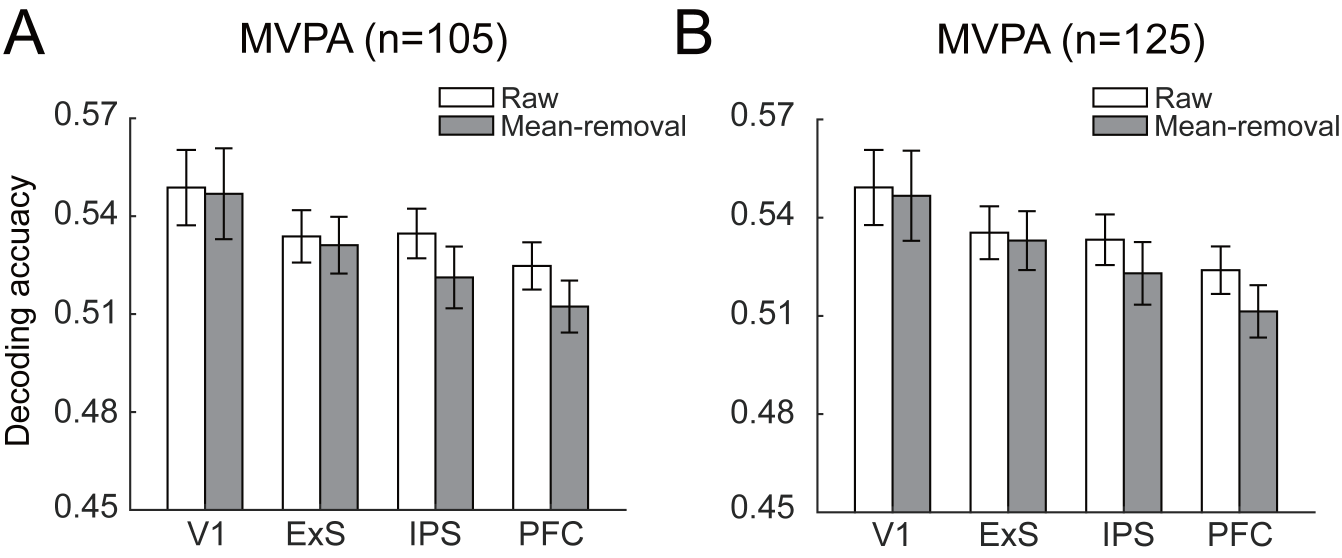

**S7 Fig. Multivariate pattern analysis before and after removing the grand mean from each attention condition.**

We evaluated the stability of MVPA decoding accuracy before and after mean removal using different number of voxels. (A) Results using 105 voxels: two-way repeated-measures ANOVA (region group × mean removal) revealed a main effect of mean removal (F_(1,141)_=8.97, *p*<0.01, η^2^=.160), and a significant interaction between region group and mean removal (F_(3,141)_=4.11, *p*<0.01, η^2^=.080). (B). Results using 125 voxels: two-way repeated-measures ANOVA (region group × mean removal) revealed a main effect of mean removal (F_(1,141)_=7.11, *p*=0.011, η^2^=.131), and a significant interaction between region group and mean removal (F_(3,141)_=3.07, *p*=0.03, η^2^=.061). Both sets of results resembled the results obtained with 85 voxels, where mean removal produced a decrease of decoding accuracy in IPS and PFC. Error bar denotes SEMs.
